# Extracting Mitochondrial Cristae Characteristics from 3D Focused Ion Beam Scanning Electron Microscopy Data

**DOI:** 10.1101/2022.11.08.515664

**Authors:** Chenhao Wang, Leif Østergaard, Stine Hasselholt, Jon Sporring

## Abstract

Mitochondria are the main suppliers of energy for cells and their bioenergetic function is regulated by *mitochondrial dynamics*: the constant changes in mitochondria size, shape, and cristae morphology to secure cell homeostasis. Although mitochondrial dysfunction is implicated in a wide range of diseases, our understanding of mitochondrial function remains limited by the complexity of inferring these spatial features from 2D electron microscopical (EM) images of intact tissue. Here, we present a semi-automatic method for segmentation and 3D reconstruction of mitochondria, cristae, and intracristal spaces based on 2D EM images of the murine hippocampus. We show that our method provides a more accurate characterization of mitochondrial ultrastructure in 3D than common 2D approaches and propose an operational index of mitochondria’s internal organization. We speculate that this tool may help increase our understanding of mitochondrial dynamics in health and disease.

## 2 Introduction

Mitochondria are the main producers of adenosine triphosphate (ATP) via oxidative phosphorylation in eukaryotic cells. Consequently, the study of their function is relevant in a plethora of diseases. While functional analysis of mitochondria can be performed on fresh, untreated tissue [5, 6] it is not applicable to preserved specimen. To this end, mitochondrial function is thought to be closely linked to its structural characteristics. Specifically, the electron transport chain is located on mitochondrial cristae (fig. 1). Accordingly, morphological features that may relate to function include (i) crista membrane (CM) surface area, which may scale with the capacity for respiratory ATP production, and (ii) crista shape, as the assembly and stability of electron transport chain complexes depend on it [7]. In particular, high local CM curvature near complex V (ATP synthase) may facilitate ATP production [8]. Dimerized complex V imposes local curvature [9, 8] and loss of dimerization results in wider cristae with blunt apices [10]. There is therefore an urgent need for methods to reliably extract critae properties.

**Figure 1:**
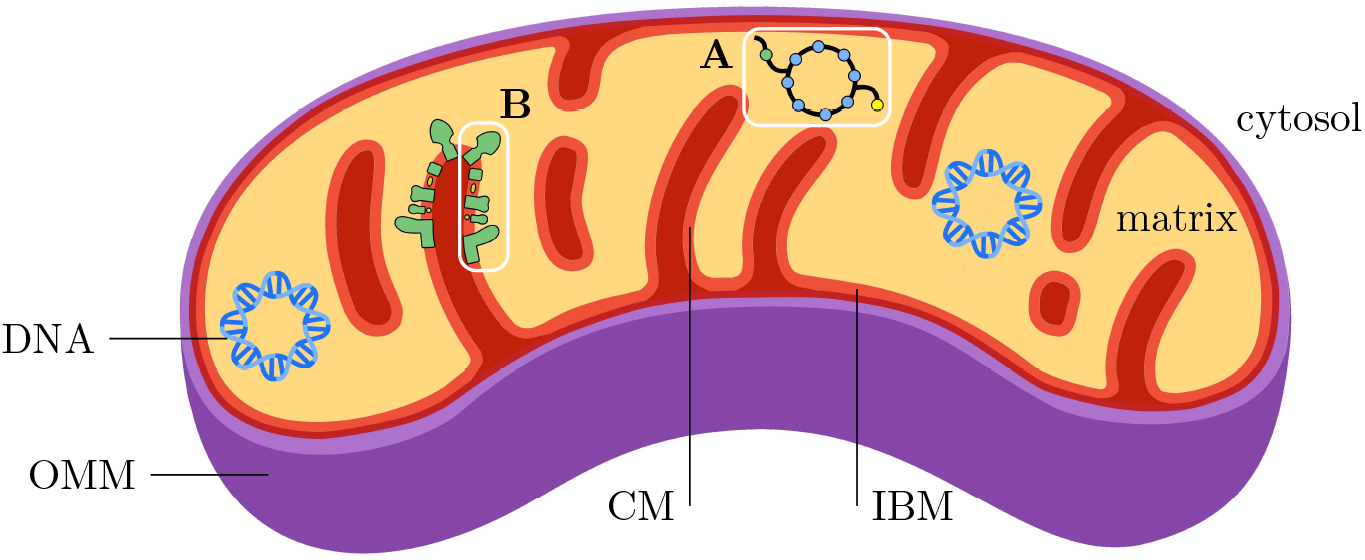
Structure of mitochondria. Mitochondria are characterized by two membranes that define three functional compartments. The outer mitochondrial membrane (OMM) acts as a barrier between the cytosol and the *intermembrane space*. The inner mitochondrial membrane (IMM), in turn, separates the intermembrane space from the *mitochondrial matrix*. In doing so, the IMM forms numerous invaginations into the matrix, the *mitochodrial cristae*. At the base of these cristae, the crista junction separates the IMM into crista membrane (CM) and inner boundary membrane (IBM), respectively, of which the latter runs largely parallel to the OMM, separated by the intermembrane space. The OMM is highly permeable to small solutes and contains proteins that allow larger molecules to pass. The IMM, however, acts as a tight diffusion barrier that only allows the passage of certain molecules via specific transport proteins. This enables the maintenance of a proton gradient between the intermembrane space and the matrix, which is critical for adenosine triphosphate (ATP) synthesis in the CM. While the matrix provides enzymes for citric acid cycle activity (A), enabling production of substrates for oxidative phosphorylation, the CM contains the respiratory chain protein complexes (B) that generate the ATP [1, 2, 3, 4].

The majority of previous and current studies demonstrating mitochondrial ultrastructure have used manual annotations on electron microscopical and -tomographical sections. This approach is labor intensive, typically restricting analyses to a limited number of mitochondria. Most often subsequent evaluation of crista proporties has been performed in 2D [11, 12, 13, 10] and the validity of generalizations to 3D is questionable. Fortunately, studies of mitochondria and cristae in 3D based on manual reconstructions of individual mitochondria have also been performed for decades [14, 15, 16, 17, 18]. Mendelsohn et al. 2021 evaluated 12 mitochondria and within this small sample the volume estimates varied by almost an order of magnitude [16] highlighting the importance of analyzing a larger population to ensure a representative sample. This becomes even more important for the dynamic mitochondrial cristae [19, 20, 21, 22].

For large scale analysis of mitochondria and cristae in 3D to be realistic, an automated approach is highly desirable. Recent advances in machine learning have made large-scale automatic segmentation of mitochondria feasible [23, 24] but until now, such automated tools have not allowed segmentation and analysis of the structures most intimately connected to mitochondrial function: the cristae. There may be several reasons for this. First, there is a scarcity of sufficiently large training datasets to train 3D segmentation models to identify cristae. Secondly, the distinction of cristae from other features in gray scale images from the electron microscope is difficult. Thirdly, well-defined distance and curvature measures in 3D, that can be extracted automatically, are needed. In this work, we suggest solutions to the challenges, enabling large scale analysis of mitochondria and their cristae in 3D based on the application of machine learning.

## 3 Material

The images used in this study were acquired by Graham Knott and Marco Cantoni from École Polytechnique Fédérale de Lausanne and are available from [25]: A 5-micrometer-cubed volume from *cornu ammonis* 1 in hippocampus from a mouse brain was imaged using Focused Ion Beam Scanning Electron Microscopy, and the mitochondria in two sub-volumes were annotated by experts. The image volume is 1065 × 2048 × 1536 voxel^3^, and the sub-volumes are 165 × 1024 × 768 voxel^3^. Voxel size is approximately 5 × 5 × 5 nanometer^3^. The original images show a small drift, hence, we performed image registration using vesicle-based drift correction [26]. As a precursor of our work, we have extended the annotations in the two provided sub-volumes to 5 classes: background, mitochondria, mitochondrial matrix, CM, and intracristal space. Examples of the data together with the initial segmentation of mitochondria are shown in fig. 2. To access the registered images and the annotated sub-volumes, please refer to section 7.

**Figure 2:**
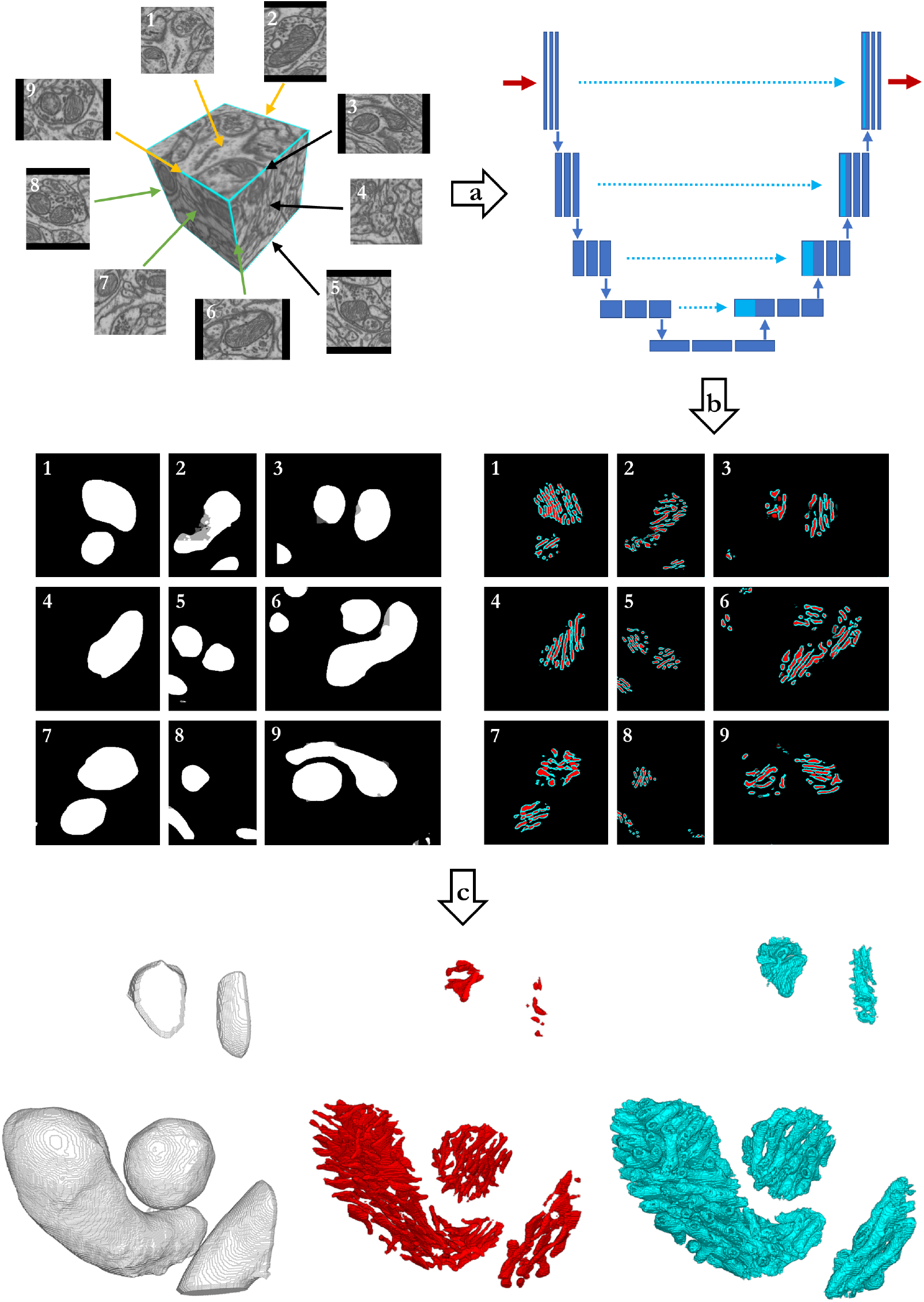
A Pipeline for segmenting cristae using the Multiplanar UNet. After drift is corrected by image registration, 2D image slices are extracted from nine different resliced planes. Selected image slices are annotated and fed to UNet for training in step (a). After training completes, the UNet is used in step (b) to segment the mitochondria (white), crista membrane (cyan), and intracristal space (red) in all nine planes. The resulting 2D stacks of segmentations from the different planes are combined using averaging, and then cleaned up by mathematical morphology in step (c) to produce the final 3D segmentation. The segmentations produced by this pipeline form the basis for all subsequent analyses.

## 4 Results

### 4.1 Multiplanar UNet overcomes the 3D training set barrier

Supervised machine learning models require correctly labelled datapoints. For 3D segmentation models, the datapoints are image sub-volumes that need to be densely annotated by experts for proper model training and validation. This is a tedious process that involves the manual annotation of hundreds of consecutive image slices for each sub-volume, possibly introducing a directional bias since the annotator tends to label slice-wise in a single slicing direction. In contrast, 2D models work on image slices, which are considerably less labour intensive to annotate, and models are often smaller and require less data to converge. However, the 3D structural information is lost.

To get the best from both the 2D- and 3D models without their weaknesses, we implemented a slightly modified version of the Multiplanar UNet [27]. This model is a 2D UNet [28] that segments a 3D volume by merging 2D segmentations of images resliced in different orientations as shown in the top left of fig. 2. At each voxel, the model produces several label-candidates as a function of reslicing-orientation, and these are merged using averaging.

We incorporated the Multiplanar UNet in an active-learning approach to manual cristae annotation to facilitate the generation of new segmentation datasets. For details, see section 6.1. The final segmentation workflow is illustrated in fig. 2.

### 4.2 Persistent homology allows distance and curvature measurements in 3D

Manual analyses of cristae and their organization poses several challenges. One is the lack of well-defined ways to measure relevant parameters. In the case of distance measurements, the choice of endpoints is not well-defined (see fig. 3) making cross-study comparisons difficult. Even if clear definitions were available, minor deviations in adherence due to limitations in manual precision could have a large effect. Manual analyses are also often affected by 2D limitations, because while many software tools allow rotations of a 3D image volume, the interactive elements are designed in 2D for selection accuracy, which may introduce a measurement bias. Furthermore, a large number of measurements is needed for statistical validity, and this is not always feasible in manual analysis.

**Figure 3:**
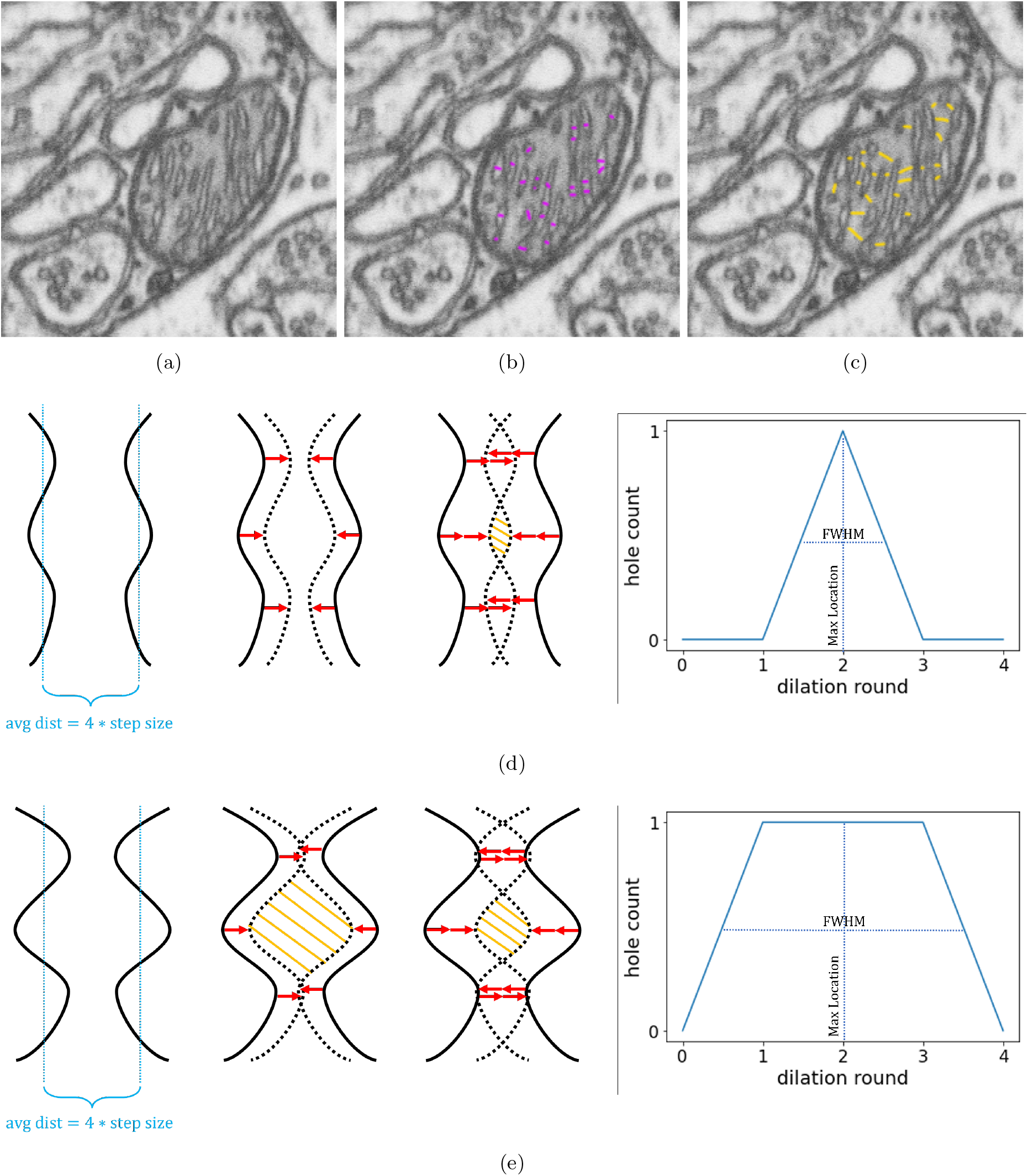
Persistent homology standardizes shortest distance measures in mitochondria. (a) – (c): Some of the many possible ways to measure crista widths (purple) and intercristal distances across matrix (yellow). (d) and (e): Persistent homology uses dilation and hole counting to standardize distance measurements. As pixels are added in dilation (red arrows), the crista membranes will move closer (dotted line) and a hole (yellow hatching) will form and eventually disappear. In crista membranes with a higher degree of curvature (e) holes will be present for more dilation rounds. This is indicated by a larger full-width-half-maximum in count curves generated during analysis, The max location in (d) and (e) remains the same at dilation round 2, which is equivalent to the average half distance between crista membranes.

As a solution to these challenges, we used the concept of persistent homology to provide a standardized and directionally unbiased way to measure cristae distances and surface curvatures in 3-dimensional space. The idea behind persistent homology [29] is to use features that exist and vary across a large parameter range to describe data that may be difficult to describe directly, since their persistence is a sign of the real signal, rather than random effects from noise, sparseness or high dimensionality. In our case, the persistent features are the count curves of holes and objects that form as a function of the number of voxels being added to or removed from our segmentation surface by mathematical morphology. Since morphology adds or removes voxels uniformly in all directions, the rate of hole or object formation depends solely on the shape of our segmentation. This means that it is possible to extract the features we need directly from the count curves, which we accomplish using the location of the maximum count (max location) and the full-width-half-maximum (FWHM). For more details on how this is done and the reasoning, please refer to section 6.2.3.

### 4.3 Gross mitochondria morphology

For the mitochondria fully contained in the Lausanne dataset, we found the mean volume to be 0.048 ± 0.059 *μm*^3^, where the uncertainty is given by the standard deviation (std). The median mitochondrial volume is lower at 0.031 ± 0.017 *μm*^3^, where the uncertainty is given as the median absolution deviation (mad). The mean mitochondrial surface area was found to be 0.86 ± 0.92 *μm*^2^, and the median mitochondrial surface area is again lower at 0.61 ± 0.30 *μm*^2^. Volume and surface area distributions are shown together with their correlation in fig. 4. The mitochondrial volume and surface area show a near-perfect linear relationship with a Pearson correlation value of 0.98 (*p* = 6.9 * 10^-258^).

**Figure 4:**
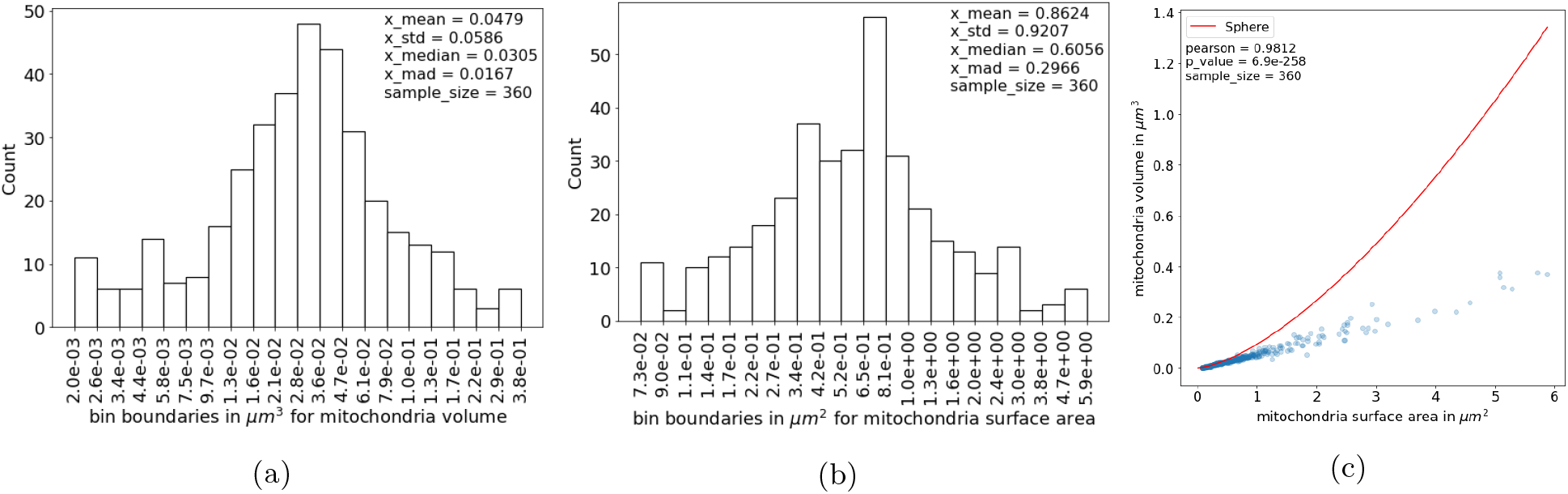
Morphology of mitochondria. The distribution of volumes (a) and surface areas (b) of mitochondria in 3D are shown for the full dataset. Please note that the counts and bin boundaries are computed on log-transformed values. The correlation between mitochondrial surface area and volume is almost perfectly linear. The curve represents the relationship between surface area and volume for a sphere (c). mad; median absolute deviation, std; standard deviation.

### 4.4 Cristae morphology and organization

The mean and median volume of the intracristal space in murine hippocampal mitochondria was found to be 3.6 * 10^6^ ± 4.8 * 10^6^ nm^3^ and 2.2 * 10^6^ ± 1.7 * 10^6^ nm^3^, respectively. The mean and median CM surface area facing the matrix side was 1.2 * 10^6^ ± 1.6 * 10^6^ nm^2^ and 7.9 * 10^5^ ± 5.1 * 10^5^ nm^2^. Volume and surface area distributions for the cristae are plotted in fig. 5.

**Figure 5:**
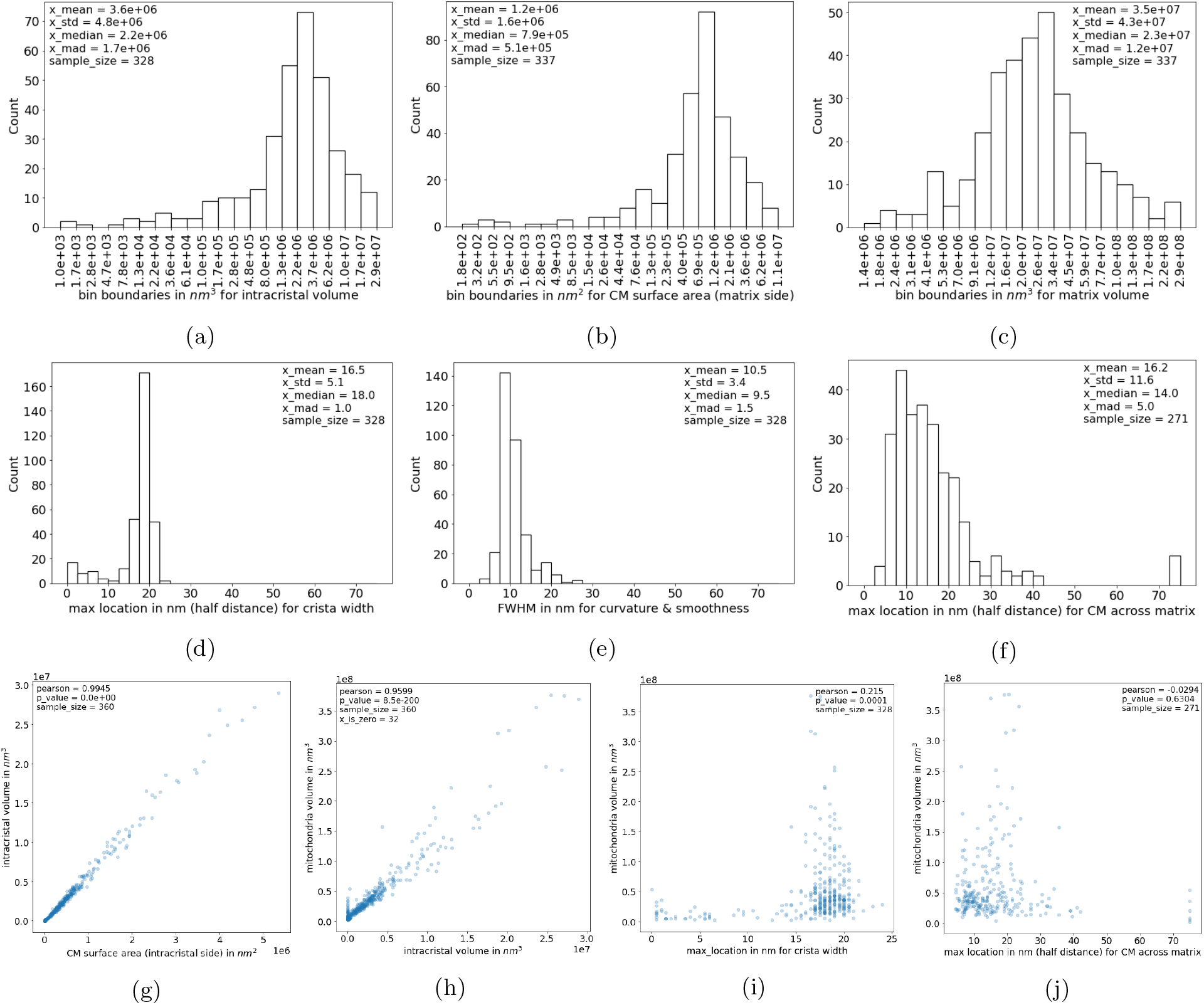
Morphology and organization of cristae. The distribution of total volumes of the intracristal space (a) and surface areas of mitochondrial cristae (b) in 3D are presented. Distributions of average crista widths (d), cristae curvature/roughness (e), and average distances between cristae across the matrix (f) in 3D is also shown. Distribution of volumes of the mitochondrial matrix is given in (c). The correlations between cristae surface area (intracristal side) and intracristal volume is almost perfectly linear (g). A positive linear relationship between the volumes of mitochondria and their intracristal spaces, respectively, is also visible (h). No clear relationship is detected between mitochondria volume and cristae width (i) or intercristal distance (j). Please note that the counts and bin boundaries in histograms are computed on log-transformed values to better visualize the data, as the untransformed data are highly left skewed. For 32 out of 360 mitochondria it was not possible to detect the inner mitochondria membrane. In sub figure (h) this results in the vertically scattered data points near the origin. Since the segmentation of these mitochondria does not seem to be erroneous, the data points are included in the scatter plot. The sample sizes vary for plots due to the differences in inclusion criteria for the parameters. CM; crista membrane, FWHM; full-width-half-maximum, mad; median absolute deviation, std; standard deviation.

The intracristal volume and the CM surface area (intracristal side) show a strong Pearson correlation of 0.99 *(p* = 0.0 * 10^0^). We measured the crista width, defined here as the minimum distance across the intracristal space (CM included), using persistent homology as described in section 6.2.3. The mean and median crista widths were 33.0±10.2*nm* and 36.0±2.0*nm*, respectively (fig. 5d). Another parameter extracted from persistent homology, the FWHM of the count curve, indirectly measures the relative smoothness and curvature of cristae. The mean and median FWHM are 10.5 ± 3.4*nm* and 9.5 ± 1.5*nm*. Generally, the smoother and less curved the CM is, the smaller the FWHM-value is, see fig. 3 and section 14.2.

The mean minimum distance between CM across the matrix is 32.4 ±23.2*nm* and the median is 28 ± 10.0*nm* (fig. 5f). This represents the average distance between individual cristae in muringe hippocampal mitochondria.

A positive linear relationship between the volume of a mitochondrion and the volume of its intracristal space was seen (Pearson correlation of 0.96 (*p* = 8.5 * 10^-200^)). The volume of the mitochondrion, however, has no clear correlations with the width of its cristae and the distance between its cristae across the matrix (fig. 5i and fig. 5j). The mitochondrial matrix has a mean and median volume of 3.5 * 10^7^ ± 4.3 * 10^7^*nm*^3^ and 2.3 * 10^7^ ± 1.2 * 10^7^*nm*^3^ (fig. 5c).

### 4.5 Consistency between 2D and 3D mitochondria measures

We evaluated the correlation between 2D and 3D shape features from mitochondria in an attempt to assess the usability of 2D measurements. When comparing measures made on a single mitochondrion in 2D, the average error with respect to the 3D measure is around 86% for the parameters evaluated here, see table 1. If a subset of 25, 50 or 100 mitochondria are measured in 2D, the average error of the mean is reduced to approximately 22%, 16 %, or 11%, respectively. Average errors for individual parameters can be found in table 1 and the results of our correlation analysis are visualized in fig. 7. The variance of the expected 3D value, which is equivalent to the square of the error, changes approximately by a factor of 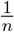, as should be expected by the law of large numbers.

**Table 1:** Correlation between 2D and 3D measurements of different parameters on a random subset of mitochondria. Presented are average percentage errors of 2D parameter means. Please refer to fig. 7 in supplemental materials to see the full error functions. n; number of mitochondria in subset

| Parameter | n = 1 | n = 25 | n = 50 | n = 100 |
| --- | --- | --- | --- | --- |
| Mitochondrial Volume | 85.01% | 22.15% | 15.71% | 11.27% |
| Mitochondrial Surface Area | 83.28% | 19.49% | 13.82% | 9.97% |
| Intracristal Volume | 86.10% | 23.20% | 17.13% | 12.06% |
| Cristae Surface Area | 88.12% | 23.01% | 15.92% | 11.49% |

**Figure 6:**
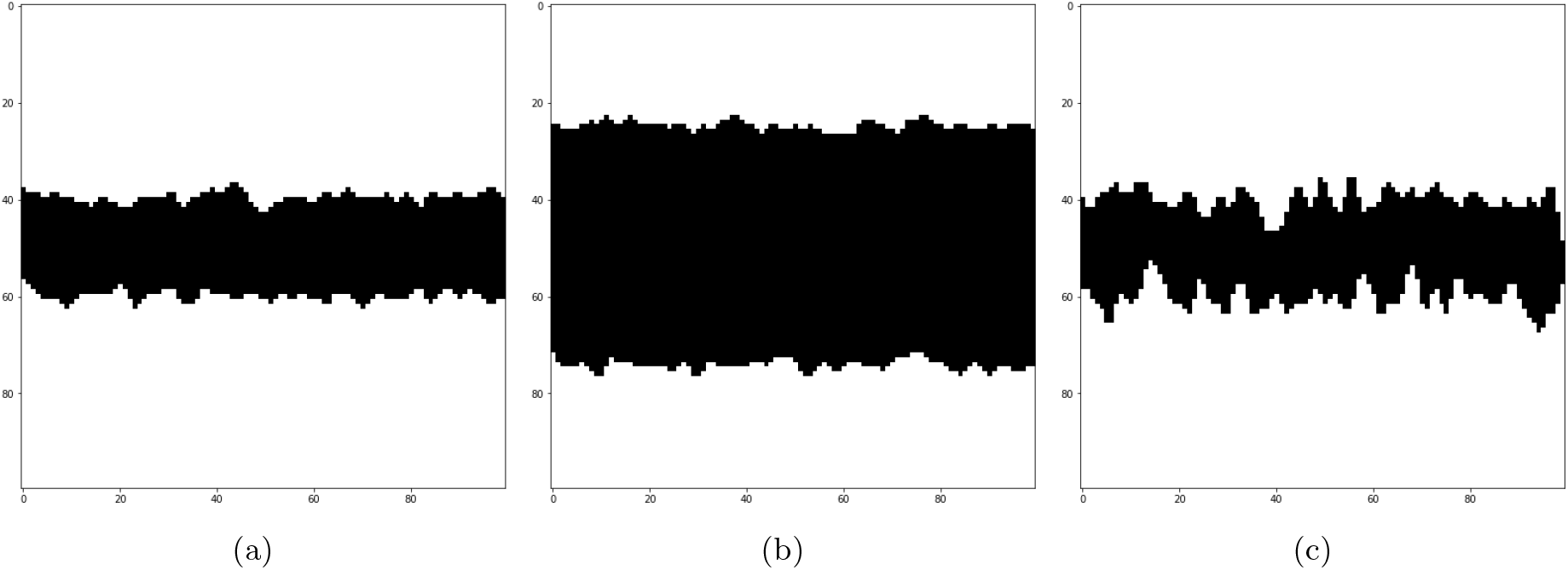
Image examples from (a) dataset 1 with *μ*_1_ = 40, *μ*_2_ = 60, *σ*_1_ = *σ*_2_ = 2, *σ*_3_ = 1, (b) dataset 2 with *μ*_1_ = 25, *μ*_2_ = 75, *σ*_1_ = *σ*_2_ = 2, *σ*_3_ = 1, (c) dataset 3 with *μ*_1_ = 40, *μ*_2_ = 60, *σ*_1_ = *σ*_2_ = 5, *σ*_3_ = 1

**Figure 7:**
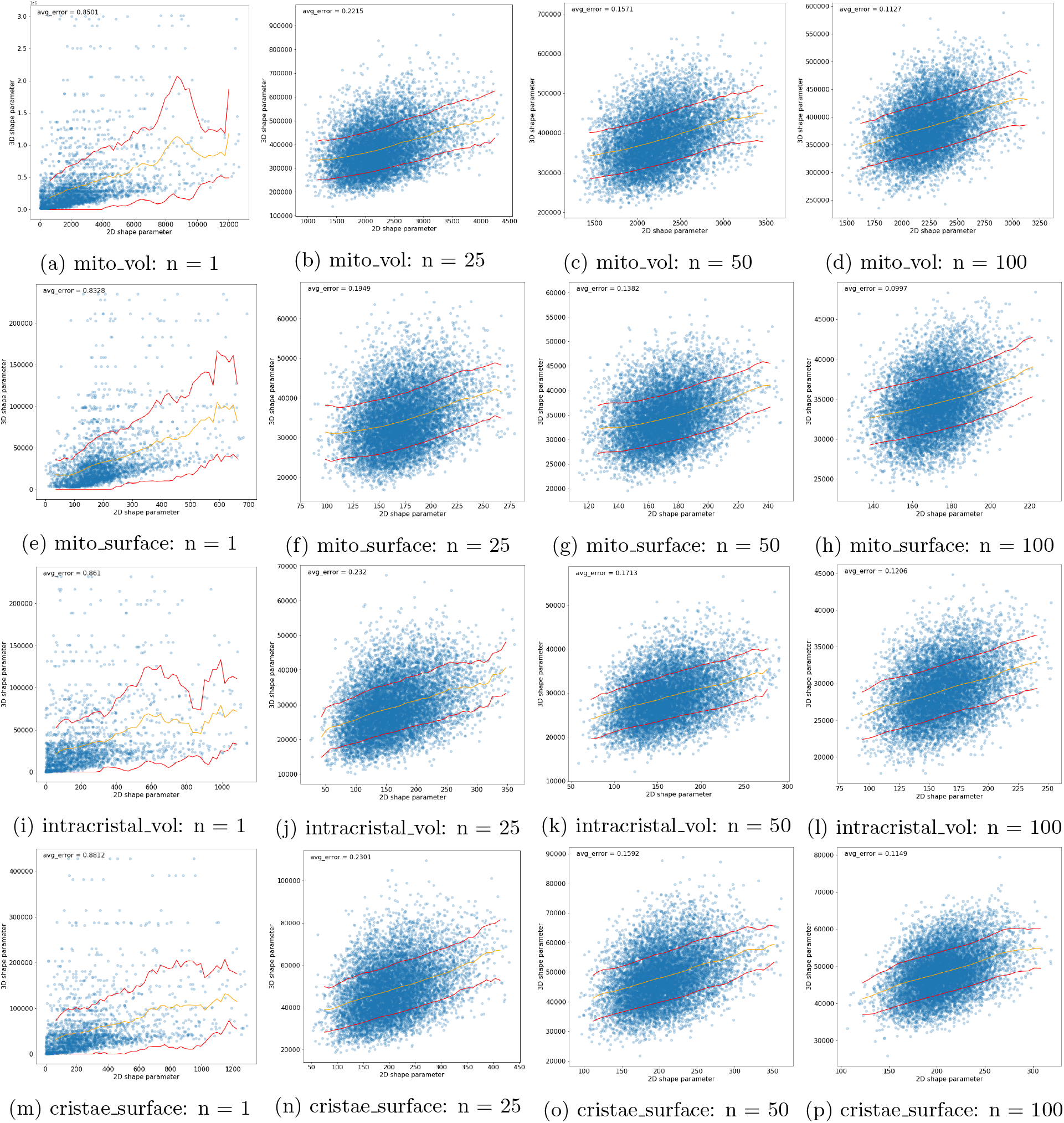
Comparison of 2D and 3D measurements. Shape parameters in 2D are measured using image slices from random planes and random slice numbers in the structure of interest, and compared against their 3D equivalents in random subsets of size n = 1 to n = 100. The comparison are done in terms of subset means and are plotted in a scatter plot. To estimate the mapping function from 2D to 3D, a sliding window is applied along the 2D value range on the x axis, where at each step, we calculate the mean and standard deviation using the 3D values contained within the window. The mean acts as the expected 3D value for a given 2D value range and the standard deviation acts as its upper and lower bound. The function curves are respectively overlaid on top of the scatter plot using yellow and red.

## 5 Discussion

We have presented a pipeline enabling semi-automatic analysis of mitochondrial ultrastructure in 3D from series of electron microscopical images. With a relatively small set of manually annotated images, automatic segmentation of the outer mitochondrial membrane, CM, and intracristal space from individual mitochondria was possible using the multiplanar 2D UNet. Subsequent estimation of 3D shape parameters for mitochondria and their cristae, along with an assessment of the distance between cristae in 3D, forms the basis for evaluation of a population of mitochondria in a tissue of interest. The described methods are directly applicable in the study of conditions affecting mitochondria.

Using persistent homology, it is possible to acquire estimates for crista width (fig. 5d) and relative curvature of the CM (fig. 5e) in 3D. Local curvature of the CM has been proposed to enable proton up-concentration near complex V, thereby increasing the possibility for production of ATP [8]. In addition to the connection between complex V dimerization and crista width [10], the width might also affect the diffusion distance for cytochrome c from the crista lumen to complex IV. Remodelling of mitochondrial cristae occurs as part of adaptive responses to altered energy substrate availability [30, 7, 31] and during apoptosis [1, 7, 32]. A cell-type independent coupling between synaptic function, CM surface area, and crista shape has also been found [17]. Of note, it should be kept in mind that cristae shape is dependent on phylogenetic group [18, 33, 34]. Cristae change their configuration dynamically through elongation or shortening, and through detachment from or fusion with the inner boundary membrane [19, 14]. They can also temporarily fuse with each other to form networks [20, 19, 21].

Persistent homology can also provide information about intercristal distance (fig. 5f). A relatively homogenous intercristal distance could be important for keeping the respiratory chain in a short distance to required substrates including oxygen and adenosine diphosphate. For the mitochondria population in a sample, the mad on the intercristal distance indicates how well-organised the cristae are. A low mad indicates that the distance between individual cristae is relatively homogenous across mitochondria whereas a high mad indicates heterogeneity. Rajab et al. 2022 applied a semi-quantitative scoring system to evaluate this [35]. This is time consuming, subjective, and requires a method for random selection of mitochondria to include in the analysis. The ratio between the intracristal volume and the mitochondrion volume also provides information about the organization of the mitochondrion. If small it indicates that the mitochondrion has lost its inner characteristics i.e. cristae. In (fig. 5h) a vertical line of data points represent these mitochondria.

In addition to the evaluation of cristae, the assessment of gross mitochondrial morphology may give us a crude indication of tissue state. Mitochondrial morphology is modulated by cycles of fusion and fission events [36] adapting the mitochondrial network to the availability of substrates and metabolic needs of the cell [30, 37]. The selective fusion of mitochondria enables transfer/sharing of organelle components [36] and allows for a more efficient energy conversion during substrate deficiency and acute oxidative stress [30, 37, 7]. Fission is involved in the removal of dysfunctional mitochondria [36] and is a main event in apoptosis [38]. After fission, some mitochondria daughter organelles are depolarized targeting them for autophagy [36]. The shape of mitochondria is furthermore crucial for their proper axonal transport and distribution [39]. An indication of overall mitochondria shape is provided by the correlation between volume and surface area. In the population of mitochondria examined here, the relationship between mitochondrial volume and surface area is linear in accordance with a previous study of mouse cerebellum [16]. A curve representing the relationship between volume and surface area for a sphere has been added to the plot (fig. 4c). *In vitro*, elongated mitochondria have been shown to be spared during autophagy while more spherical mitochondria were not [30]. If the points in the correlation plot fall on the curve for a sphere, a plausible guess would therefore be that the mitochondria are damaged. The curve may also help in the evaluation of data reliability. A sphere has the smallest possible volume to surface area ratio so no points in the correlation plot should fall above the curve.

In 2D, the results of morphometric analyses depend on characteristics of the region of interest. If rotational invariance is present in the tissue, meaning that the distribution of size, orientation, and shape of objects is independent of the direction of imaging planes, then valid information can be obtained if enough images are analysed. For many biological tissues, however, this is not the case. Our experiments on the comparison of 2D shape parameters of mitochondria against their 3D counterparts showed the importance of using 3D shape for analysis (table 1). For a subset of 100 mitochondria the parameters evaluated here can be estimated in 2D with an average error of around 11%. This means that for group comparisons, the mean difference between groups needs to be least 22% to be detectable from 2D images. Whether this is sufficient depends on the study, but for most cases, an error of 11% is too large to be useful. Further increasing the subset of mitochondria analysed will continue to decrease the error following a power law function. Evaluating the distributions of shape parameters for mitochondria estimated in 3D in this study, the majority have a mean that is significantly higher than the median. This suggests that there are a few extreme outliers affecting the mean but due to the considerable sample size, made feasible with semi-automatic detection, it is possible to identify them as outliers.

The 2D multiplanar UNet, we have used here, has the advantage of needing only a limited amount of manually annotated images to train. It has a performance (section 14.1) comparable to current state-of-the art 3D models for segmenting mitochondria [40] and is less labor intensive. Gross mitochondrial shape parameters in rodent brain [17, 23, 16], and crista width based on manual reconstruction in rat- and chick nervous tissue, and Hela cells [14, 15, 19], have previously been estimated in 3D. Our results (fig. 4, fig. 5d) are in line with the previous findings. Our measure of intercristal distance (fig. 5f) deviates from a previous finding in Hela cells (2D) [41]. It is unknown if the discrepancy stems from species- and tissue variation, methodological differences, or 2D to 3D discrepancy.

The internal organization of mitochondria is increasingly being evaluated, and focus on mitochondrial cristae changes in the study of disease has been seen in different research fields [42, 11, 35]. Cristae have been shown to increase in width and have more rounded apices under hypoxic conditions *in vitro* [43] and in patients with the oxidative phosphorylation disease Leigh syndrome [10]. Gomes et al. 2011 observed a connection between elongation of mitochondria and increased cristae density during starvation (2D, *in vitro* and *in vivo*) [30], altered organisation of cristae have been shown in ovarian cancer (2D, *in vitro*) [44] and after cerebral ischemia (2D, *in vitro)* [11], and ischemic stroke resulted in a loose, heterogeneic organization of cristae (2D, *in vivo*) [11]. The changes are most likely related to the pathological conditions as combining results about gross mitochondrial volume with information about internal distances in our sample suggests that the organization of the inner mitochondrial membrane is independent of mitochondria size (fig. 5i, j). A pattern of cristae of comparable width is merely repeated more times in large mitochondria. Additional experiments are needed to determine if this is general for normal tissue. Even though there may be challenges with interpretation of 2D analyses, the results presented here indicate the relevance of examining a spectrum of mitochondria of different sizes in disease and this is feasible with the methods described here.

A limitation to our analyses is that only mitochondria, where it is possible to segment cristae, can be included. The subpopulation where this is not possible may be of poorer quality due to organelle degradation, it may be an issue related to tissue processing, or a combination of the two. To evaluate the potential impact of this undesired selection, we suggest always assessing the fraction of the whole these mitochondria constitute. In our study 9% of the evaluated mitochodnria were of this type. An additional challenge is that mitochondria gross shape parameters are affected by cellular location [24, 14, 45]. Separating mitochondria into subpopulations depending on location for example in the cell soma or in processes require larger 3D volumes with concomitant increased imaging times and data amounts. Thirdly, individual mitochondria with touching membranes may be segmented as one. This potentially complicates the distinction between individual mitochondria and an interconnected network. However, with increasing sample size the impact of this error decreases. Moreover, it is always possible to go back and evaluate the raw images. To strengthen the results from ultrastructural analyses, they can be combined with an evaluation of mitochondrial function ex. via an analysis of glycolytic and aerobic metabolism [5]. Changes in ultrastructure may also be evaluated further using molecular biological analyses of important components in cristae assembly.

In conclusion, we provide a method for detailed analysis of mitochondrial ultrastructure in 3D based on a deep learning algorithm. From a limited amount of images with manually annotated ground truth data, we were able to reliably segment the mitochondria and their cristae. From these segmentations, we extracted information about crista surface area, volume, and shape. Furthermore, with the persistant homology method, introduced in this article, we derived statistical summary information about the internal organisation of the cristae. Our new method is not restricted to cristae structures but can be applied to any other tubular shape. Future work will include an in depth investigation of its applicability to other shapes of biological relevance.

## 6 Methods

In this work, we have employed a standard deep-neural network for segmenting the images, estimated standard geometric object features, used a topology measure to characterize long-range object relation, and investigated the relationship between parameters estimated from 2D slices and measured in 3D, respectively. All of this will be detailed in the following.

### 6.1 Image Segmentation with the Multiplanar UNet

All segmentations in this study were done using a modified version of the Multiplanar UNet [27]. The idea is that by merging 2D segmentation results from multiple planes at different angles, we can compensate for the loss of 3D information in 2D, because structures not seen in one plane will most likely be visible in another plane. However, since the core segmentation model is 2D, we can avoid having to annotate the groundtruth masks in 3D, which is a labour intensive task. To maximize the field of view for better 3D coverage, while minimizing the number of planes for computational efficiency, we used nine planes angled at 45 degrees to each other (fig. 2).

We extended the original annotation of mitochondria in the 2 sub-volumes supplied by [25] with our own annotation of the CM and intracristal space. We used an active learning approach, iterating the following steps to build up a dataset of 150 image slices:

1. Select a few 256 × 256 pixel image slices from the image volume. In the first iteration, this would be random slices from random planes. In all subsequent iterations, the slices can either be random (like the first iteration), or be chosen from places where the model didn’t work very well.
2. Manually label or correct the CM in the cyan channel and the intracristal space enclosed by them in the red channel.
3. Train a multi-class 2D UNet, initialized with weights from the mitochondria segmentation task for transfer learning.
4. Apply the multi-class 2D UNet in a multi-planar fashion to acquire 3D results.
5. Repeat all steps until segmentation quality is sufficiently good.

The final model for CM and intracristal space was trained on all 150 image slices from above, where 80% was used for training and 20% for validation. The model was initialized with weights from mitochondria segmentation for transfer learning. Image augmentation consisting of translation, rotation, shearing, intensity adjustment and noise introduction was actively applied on the training set to compensate for the still limited amount of data. The validation set was also augmented, but only initially to ensure the same set of validation images were used in each epoch, and it excluded intensity and noise augmentations to ensure we only validate with realistic images. Model optimization was done using the ADAM optimizer on Intersection over Union (IOU) loss. This is important because of the extremely unbalanced class ratio in favour of the background. To achieve efficient training speed and optimize performance, adaptive learning rate was used, where we reduced the learning rate by a factor of 3 if the validation loss did not improve for 5 epochs, starting with learning rate = 0.0001. The training was stopped early if the validation loss did not decrease for 25 epochs.

Mitochondria toucing the boundary of the image volume, and mitochondria with a volume lower than 25^3^ voxel^3^, were excluded from the analysis. The performance of our segmentation model were evaluated using 5-fold cross-validation (see section 14.1).

### 6.2 Computation of shape measures

All shape measures are computed on a mitochondrion by mitochondrion basis and the results are collected as distributions of values.

To combine individual mitochondria with their respective CM and intracristal spaces, the mask from connected-component analysis on the mitochondria segmentation was multiplied with the segmentation of CM and intracristal space, respectivey.

#### 6.2.1 Volume

The volume is a measure of object size. Since our segmentation results are binary images, where the foreground is 1 and background is 0, the volume is a summation of every voxel value (assuming the object has been isolated). The volume unit can be converted to real-world measurement by multiplying with the image resolution (here 125nm^3^).

In this study, we measure the mitochondrion volume, the intracristal space volume and the matrix volume, the latter of which is defined as:

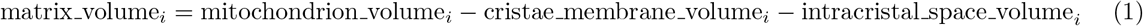

#### 6.2.2 Surface Area

To calculate the surface area of an object, the segmentation volume was initially converted to a triangular mesh using marching cubes. The output of the algorithm contains a list of all vertices (with coordinate values) and all the triangular faces (made up by the vertex indices). The total surface area of the object is then the sum of surface areas of the faces:

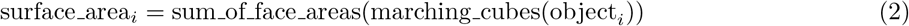

where *i* refers the *i*th object or surface area.

In this study, we measure the surface areas of the mitochondria, the intracristal space, and the CM (facing the matrix).

#### 6.2.3 Persistent Homology

In this paper, we combine persistent homology and image analysis. By applying mathematical morphology to our segmentation masks and counting the number of holes or objects that appear and disappear, we use persistent homology as an efficient and accurate way to indirectly measure object distances and relative curvatures in 3-dimensional space.

Existing implementations of mathematical-morphology-based persistent homology do exist but they are primarily designed to describe the distribution of multi-dimensional point clouds. As a result, they do not work efficiently on images and their output is structured in a way that is unsuitable for extracting the features we need. We therefore implement a custom approach to persistent homology.

The two main types of morphological operations used in our persistent homology analysis are dilation and erosion, and they can respectively enlarge or shrink an object uniformly in all directions by adding or removing a layer of voxels from the object’s surface. To enable measurements at subpixel accuracy, we took a PDE-approach to mathematical morphology. Starting with the 2D first-order Osher–Sethian upwind scheme [46, 47, 48, 49], we adapt them to 3D by introducing new terms into the square root and subsequently enforcing a value constraint between 0 and 1:

Dilation

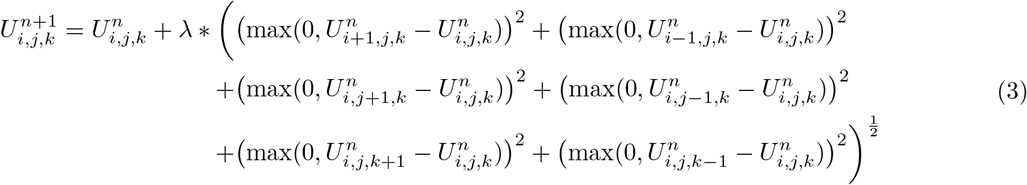

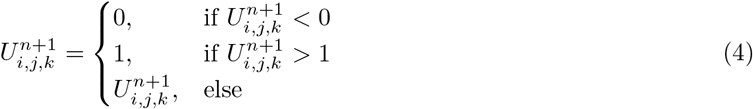

Erosion

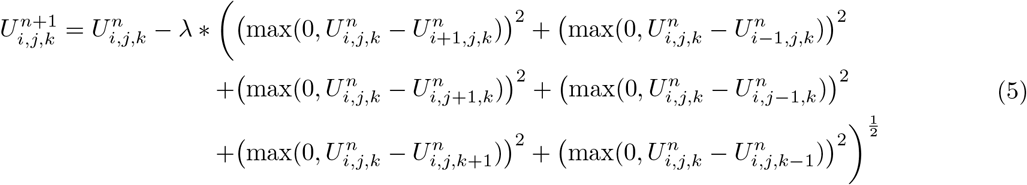

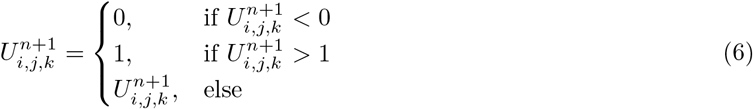

Where *U^n^* represents the segmentation volume at current timestep, *U*^n+1^ represents the segmentation volume after a single round of subpixel morphology is applied to *U^n^*, and *λ* represents the timestep. In our case, *λ* = 0.1 is used. This means that 10 rounds of subpixel morphology will be equivalent to one full round of standard mathematical morphology, and it represents the point where a single layer of voxels are either added or removed from the segmentation surfaces (depending on whether it is dilation or erosion). We apply persistent homology on a mitochondrion-by-mitochondrion basis.

Conceptually, in the case of dilation: As segmented objects are enlarged, previously non-touching points on different objects or different branches of the same object will eventually make contact with each other. When this happens, holes will begin to form in the background region and the hole count increases, see fig. 3. As the dilation continues, different objects or different branches of the same object will fully merge and the holes that formed earlier will disappear. The resulting count curve therefore acts as an indirect shape descriptor from which information can be extracted. The equivalent curve can be obtained by erosion and object counting.

To measure the **average minimum distance between cristae across the matrix**, we perform dilation and hole counting on the sum of intracristal space- and CM segmentations (e.g. red objects + cyan objects in fig. 2). For **crista width**, we also use the sum of intracristal space- and CM segmentation, but perform erosion and object counting instead.

In all cases, each round of subpixel morphology produces a non-binary grayscale mask valued between 0 and 1. Although this raw mask is always the one to be used for the next round of subpixel morphology, a binarized version with a threshold at 0.5 is needed for the counting step. For hole counting after dilation, we first multiply the binarized dilation mask with the mitochondrion segmentation to ensure the dilation does not go out of bounds. The dilation mask is then inverted, before a 3D connected components with a connectivity of 26 is performed to calculate the number of holes. Since an extra hole will always exist in the background, we subtract 1 from the resulting count. Object counting after erosion works the same way, except we go directly to counting without multiplying and inverting the mask, and we do not subtract anything from the object count.

We summarize our curves using the max location and FWHM, as measured by the iteration number. Given that the data is discrete and has a degree of randomness, the raw count curves will be slightly jagged and should be filtered using a Gaussian kernel. Due to noise susceptibility, the initial five rounds of subpixel morphology, corresponding to half of a full round of dilation/erosion, are not included when finding the max locations and their FWHM.

The max location can be interpreted as half of the average minimum distance across the region being dilated or eroded, because an iteration having the most holes/objects implies that it is also the iteration where most surface points make contact. The FWHM measures the surface smoothness and curvature of the same region. In this case, smoothness and curvature, respectively, refer to the degree of roughness and the extend to which the surface bends, but despite their slightly different definitions, surface smoothness and surface curvature are in fact the same shape parameter but on different scales. With rougher and more curved surfaces, the existence of more convexities and concavities will make holes and objects appear earlier and prolong the number of iterations it takes for them to disappear (as a result of a full merge). See fig. 3 for an example. See also supplementary material section 14.2 for a synthetic experiment illustrating the relation between our suggested measures and shapes.

To ensure numeric validity, the following filtering criteria were applied When performing statistics on the calculated max locations and FWHMs. For the average minimum distance between cristae across the matrix, we require the presence of cristae membrane and that there must exist different branches, folds, or instances of cristae membrane to measure against. In technical terms, we only include results computed from count curves with a maximum hole count ≥ 1. For crista width, we require that both the intracristal space volume and the cristae membrane volume to be larger than 0.

### 6.3 2D to 3D relationships

Initially, the connected components of our mitochondria segmentation are used to extract the 3D sub-volume for each mitochondrion. Random image slices are then sampled from the nine predefined planes orientated at 45 degrees from each other to maximize data variability, see fig. 2. The 2D perimeters and 2D crosssectional areas of mitochondria are subsequently computed from the image slices and paired against their corresponding 3D surface areas and 3D volumes for further analysis. The comparison is made for varying sample sizes ranging from n = 1, which is equivalent to a direct single-datapoint comparison, to n = 100, where a comparison is made between the 2D and 3D averages of 100 datapoints. To ensure statistical reliability, 10000 subsets of matching datapoints are randomly generated for each sample size tested.

Given a series of 2D parametric values and their 3D equivalents, we can estimate the mapping function from 2D to 3D by first plotting the 2D values against the 3D values on a scatter plot, and then sliding a window of size *ω* over the 2D value range on the x axis. For each step of the moving window, we calculate the mean and std using the 3D values contained within the window. The mean is the expected 3D value for a given 2D value range and the std its corresponding upper and lower bound: mean ± STD. Since the parameters cannot be negative, all negative lower bounds are set to 0. To eliminate noise caused by data sparsity at the higher end of 2D values, we terminate the sliding window when it contains less than 10 data points. The size *ω* should be adjusted based on the scale of the 2D values, and we found empirically that *ω* = 0.025 * max(2D_values) works well.

The degree of correlation between 2D and 3D shape parameters can be summarized by computing the average error for approximated mapping functions:

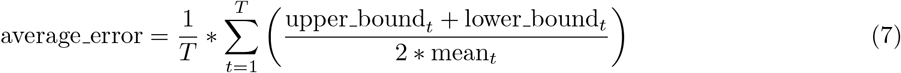

where *T* is the total number of windows and *t* is the window index.

## 7 Data availability

The Lausanne dataset is available from the website of École polytechnique fédérale de Lausanne (EPFL): https://www.epfl.ch/labs/cvlab/data/data-em/

Our cristae annotations can be found at: erda.dk.

## 8 Code availability

The code for this paper can be found at: qim.dk.

## 10 Acknowledgement

We would like to thank Peter Bross, and Jens Randel Nyengaard for fruitful discussions during this work. Vibe Sporring is thanked for technical assistance with graphical illustrations.

## 12 Author information

All the authors took part in writing the paper. The authors jointly developed the methods. C.W. wrote the code for the pipeline and ran the computer analyses. All authors reviewed the final paper.

## 13 Ethics declarations

The authors have no competing interests in this work.

## 14 Supplementary material

### 14.1 Segmentation Performance

The mathematical definitions for the performance metrics used in table 2 are provided below:

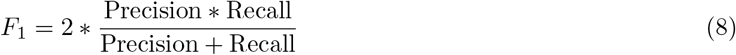

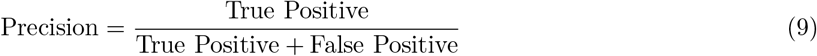

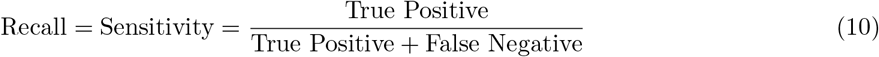

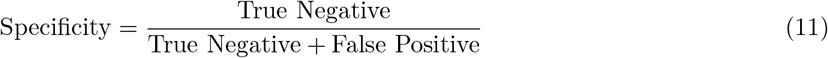

**Table 2:** Performance of the segmentation model used in this study. For the mitochondria, a 3D segmentation was produced by the 2D multiplanar UNet and then compared against the 3D groundtruth. The segmentations of crista membrane and instacristal space are evaluated in 2D (without the multi-planar aspect), because the 3D volume is not fully annotated for cristae and intracristal space.

| Parameter | F1 score (%) | Sensitivity (%) | Specificity (%) |
| --- | --- | --- | --- |
| Mitochondrion | 0.923 | 0.945 | 0.994 |
| Crista membrane | 0.578 | 0.687 | 0.991 |
| Intracristal space | 0.652 | 0.626 | 0.996 |

### 14.2 Simplified Persistent Homology Example

To demonstrate how the max count location and FWHM can be used as shape measures, we generate sample images of size *(n, m)* in 2D using the equation:

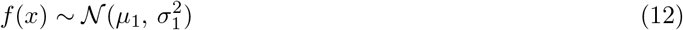

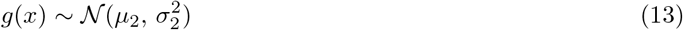

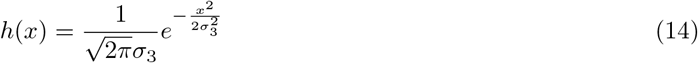

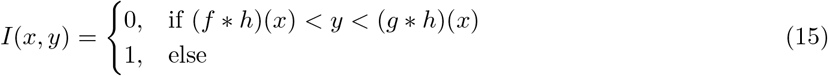

The idea here is to use 2 normal distributed and smoothed sequences, *f* * *h* and *g* * *h*, to represent neighbouring cristae membranes that we would like to measure. In our equation, *μ*_1_ and *μ*_2_ are the means of the Gaussian distribution (where *μ*_1_ < *μ*_2_), and they correspond to the theoretical row-wise locations of the membranes within our sample image I; *σ*_1_ and *σ*_2_ are the standard deviations of each sequence, which specifies the degree of roughness/curvature of the generated membranes; and finally, *σ*_3_ specifies the strength of the Gaussian smoothing filter, which is needed to reduce excessive local raggedness.

In our experiment, we generated 3 datasets of 1000 images, where each dataset has their own unique theoretical distance (*μ*_1_ – *μ*_2_) between the membranes, and unique levels of roughness/curvature (see fig. 6). As the region which we would like to measure is black in the images, the dilation and hole counting version of persistent homology is used. The measured results are shown in table 3. We see that the max location correlates with the distance between the synthetic membranes, and the FWHM correlates with the roughness of the membrane surfaces.

**Table 3:** Persistent homology results on artificially generated datasets from fig. 6. Column 1 shows the true half distances to be compared directly against the measured half distances/max locations in column 3. These generally correlate quite well, but there is a small degree of underestimation in the half distances. Column 2 records the standard deviations used for generating the curves, and is to be used for indirect comparisons against measured FWHM in column 4. The values in columns 2 and 4 are not supposed to match, but they should have the same in-column ordering in terms of value size, which is the case here. FWHM; full-widthhalf-maximum

| Experiment | $(\mu_2 - \mu_1)/2$ | $\sigma_1 = \sigma_2$ | max location | FWHM |
| --- | --- | --- | --- | --- |
| 1 | 10 | 2 | 9.43 | 1.79 |
| 2 | 25 | 2 | 24.41 | 1.77 |
| 3 | 10 | 5 | 8.13 | 3.02 |

### 14.3 2D to 3D comparison

